# Autoimmunity associated allele of tyrosine phosphatase gene *PTPN22* enhances anti-viral immunity

**DOI:** 10.1101/2023.06.28.546768

**Authors:** Robin C. Orozco, Kristi Marquardt, Isaphorn Pratumchai, Kerri Mowen, Alain Domissy, John R. Teijaro, Linda A. Sherman

**Author notes:** co-corresponding authors Direct all inquiries about this paper to Robin C. Orozco (email-; mailing address: 1200 Sunnyside Ave, Haworth Hall, Lawrence, KS 66044) or Linda A. Sherman (email-).

## Abstract

The 1858C>T allele of the tyrosine phosphatase *PTPN22* is present in 5-10% of the North American population and is strongly associated with numerous autoimmune diseases. Although research has been done to define how this allele potentiates autoimmunity, the influence *PTPN22* and its pro-autoimmune allele has in anti-viral immunity remains poorly defined. Here, we use single cell RNA-sequencing and functional studies to interrogate the impact of this pro-autoimmune allele on anti-viral immunity during Lymphocytic Choriomeningitis Virus clone 13 (LCMV-cl13) infection. Mice homozygous for this allele (PEP- 619WW) clear the LCMV-cl13 virus whereas wildtype (PEP-WT) mice cannot. This is associated with enhanced anti-viral CD4 T cell responses and a more immunostimulatory CD8α^-^ cDC phenotype. Adoptive transfer studies demonstrated that PEP-619WW enhanced anti-viral CD4 T cell function through virus-specific CD4 T cell intrinsic and extrinsic mechanisms. Taken together, our data show that the pro-autoimmune allele of *Ptpn22* drives a beneficial anti-viral immune response thereby preventing what is normally a chronic virus infection.

**Author Summary:** *PTPN22* and its alternative allele, 1858C>T, has largely been studied in the context of autoimmunity. Through these studies, researchers defined roles for PTPN22 in regulating T lymphocyte activation, myeloid cell cytokine production, and macrophage polarization. Despite these immune pathways being critical for anti-viral immunity, little work has studied how this allele impacts virus infection. In this study, we examine gene expression and function of immune cell subsets to demonstrate how a common allelic variant of *PTPN22,* which strongly increases the risk of autoimmune disease, promotes successful clearance of an otherwise chronic viral infection.

## Introduction

Allelic variation in genes that regulate immune cell responses potentially impact an individual’s response to self and foreign antigens. Genome wide association studies (GWAS) have identified variants in immune-related genes that are being increasingly associated with protective or pathologic consequences during disease [1]. However, the underlying mechanism(s) through which these mutations impact disease often remain incompletely defined.

The 1858C>T allele of the tyrosine phosphatase *PTPN22 (*causing amino acid substitution R620W) is present in 5-10% of the North American population and is strongly associated with numerous autoimmune diseases, including Type I Diabetes (T1D), rheumatoid arthritis, systemic lupus erythematosus, and others [2–10]. This alternative allele of *PTPN22* is considered the highest non-HLA risk allele in autoimmunity [7, 9]. This pro-autoimmune allele is known to affect innate and adaptive immune functions, including, lymphocyte activation, toll-like receptor signaling, and cytokine production in various autoimmune contexts [8, 11–17] [18, 19]. In humans, *PTPN22* encodes the protein Lyp that is expressed in all immune cells [10]. To study its function in immune cell types, researchers have often employed *Ptpn22* knock-out mice, which are deficient in expression of the Lyp ortholog, PEP (PEP-null) [20]. In lymphocytes, Lyp/PEP tempers T cell receptor (TCR) and B cell receptor (BCR) signaling through dephosphorylation of Src kinases. Binding partners enabling such activity include TRAF3 and CSK [12, 14]. In myeloid cells, the interaction of Lyp/PEP with TRAF3 promotes TLR activation and type I interferon production [11, 15]. Through these studies, researchers have identified mechanisms that may contribute to the pathogenesis of multiple autoimmune disorders. Despite the importance of these same immune regulatory factors in shaping a robust anti-viral immune response, the impact of the pro-autoimmune allelic variant of *PTPN22* on anti-viral immunity has received far less attention.

One of the best-defined experimental models of persistent virus infection is the lymphocytic choriomeningitis virus clone 13 (LCMV-cl13) model in C57BL/6 mice [21–24]. Using LCMV clone 13 (LCMV-cl13) researchers have defined mechanisms of viral persistence and immune cell exhaustion [24–27]. Previously we and others found that, upon infection with LCMV-cl13, mice lacking *Ptpn22* (PEP-null) have accelerated viral clearance and exhibit less of an exhausted T cell phenotype [28, 29]. Specifically, there was enhanced anti-viral CD4 T cell function and improved CD8 T cell function late in infection, suggesting *Ptpn22* contributes to the generation of T cell exhaustion. However, these studies were performed using PEP-null mice rather than mice expressing the equivalent of the human pro-autoimmune allele. Therefore, it remained to be determined whether the alternative human allele would similarly contribute to viral clearance and immune effector functions.

In this study, we use C57BL/6 mice mutated using CRISPR/Cas9 to express the murine ortholog of the *Ptpn22* pro-autoimmune allele (PEP-619WW) and LCMV- cl13 to define new mechanisms by which the autoimmunity associated allele of *PTPN22* contributes to viral clearance and enhances anti-viral T cell and myeloid cell responses.

## Results

### Ptpn22 alternative allele promotes LCMV-cl13 viral clearance

Mice expressing the *Ptpn22* wild type allele (PEP-WT) or *Ptpn22* pro autoimmune allele (PEP-619WW) were infected with LCMV-cl13. All mice lost weight within the first week of infection (Figure 1A). However, PEP-619WW mice stabilized and regained their original weight more quickly than PEP-WT mice (Figure 1A). This difference also correlated with viral titers in the serum of these animals (Figure 1B). While PEP-WT mice had detectable virus in their serum out to day 41, viral titer in PEP-619WW mice decreased by 22 days post infection (dpi) (Figure 1B). PEP-619WW did not restrict early LCMV-cl13 infection, as viral loads in both genotypes of mice were high 3 dpi in sera, lungs, livers, and kidneys (Figure 1B-C). By day 41, PEP-619WW mice had no detectable virus in the serum and largely cleared virus from the target organs spleen, lungs, and livers (Figure 1D). In contrast, PEP-WT mice still had detectable virus in these organs. In both strains of mice, virus was still detectable in the kidney, an organ in which it is known virus remains detectable despite clearance from other organs [22, 23, 30, 31].

**Figure 1.**
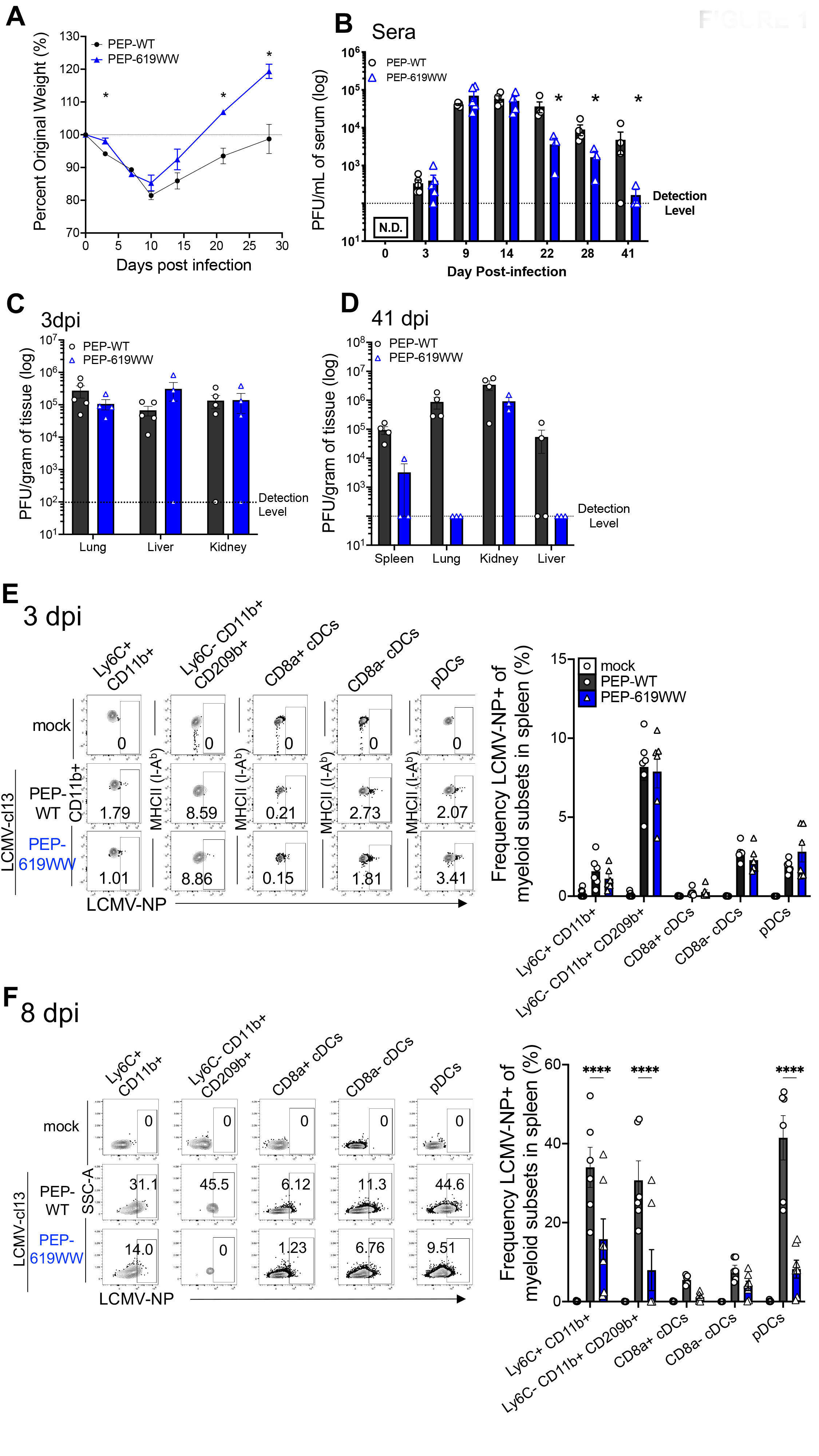
PEP-619WW mice clear LCMV-cl13 infection. C57BL/6 mice with wild type (PEP-WT, black) or pro autoimmune allele (PEP-619WW, blue) *Ptpn22* gene were infected with 1x10^6 PFU chronic viral strain LCMV-clone 13 (LCMV- cl13) (i.v). On indicated days mice were weighed (A), and bled for serum titer (B). Viral load was determined at early (3dpi, D) and late (41dpi, E) within target organs. Using flow cytometry presence of LCMV nucleoprotein (LCMV-NP) (clone VL-4) was determined in multiple myeloid cell subsets at day 3 (E) and day 8 (F). Representative flow plots (which is the median mouse data point in corresponding bar graph) shows LCMV-NP+ cells for different splenic myeloid subset at each time timepoint for mock infected (white bar, circle), infected PEP- WT (black bar, circle), and infected PEP-619WW mice (blue bar, triangles) Quantification for Frequency of LCMV-NP+ cells in corresponding bar graph below. Gating strategy for all myeloid subsets is: Lymphocytes> Single cell x2> Live> CD3- CD19-> Ly6G- CD11b+/-> (non-neutrophils). Ly6C+ CD11b+ cells are: myeloid cells> F4/80+ CD11c-> Ly6C+ CD11+. Ly6C- CD11b+ CD209b+: myeloid cells> F4/80+ CD11c-> Ly6C- CD11b+> CD209b+. CD8α+ cDCs: myeloid cells> F4/80- CD11c+> PDCA-1- CD8α+> MHC-II (I-Ab)+. CD8α- cDCs: myeloid cells> F4/80- CD11c+> PDCA-1- CD8α-> MHC-II (I-Ab)+. pDCs: myeloid cells> F4/80- CD11c+/-> PDCA-1+ CD8a+/- > Ly6C+ B220+. Weight loss studies pooled from 3 separate experiments: PEP-WT n=15, PEP-619WW n=20. Serum titer, tissue titer from representative experiment. Each dot indicates an individual mouse. Quantification of LCMV-NP protein in myeloid subset is from representative experiment per day, each dot represents an individual mouse. SEM shown. *p<0.05, **p<0.01, ***p<0.001, ****p<0.0001, Two Way ANOVA with Sidak Post Hoc Analysis.

Cellular tropism is thought to be a key factor in determining the potential chronicity of a virus infection [25, 32]. Specifically, the ability to establish a persistent infection in C57BL/6 mice is attributed to the increased capacity of LCMV- cl13 to infect dendritic cells (DCs) and macrophages, including plasmacytoid DCs (pDCs) and marginal zone macrophages [25, 32–34]. To determine if PEP-619WW mice had altered tropism we examined LCMV nucleoprotein (NP) expression in multiple myeloid cell subsets. At 3 dpi, no difference was detected in LCMV-NP+ cells amongst monocytes (Ly6C+ CD11b+), marginal zone macrophages (Ly6C- CD11b+ CD209b+), CD8α^+^ conventional dendritic cells (cDCs), CD8α^-^ cDCs, or pDCs (Figure 1E). By 8 dpi, PEP-619WW mice had a significantly decreased proportion of LCMV-NP+ splenic monocytes, marginal zone macrophages, and pDCs (Figure 1F). There was also a lower proportion of CD8α^+^ cDCs that were LCMV-NP+ cells, but this did not reach significance. We did not observe any difference in the proportion of LCMV-NP+ CD8α^-^ cDCs (Figure 1F). At 8 dpi, the marginal zone macrophages from most PEP-619WW mice had no detectable virus (Figure 1F). However, in 2 PEP-619WW animals, about 20-30% of marginal zone macrophages were LCMV-NP+ (Figure 1F). This is in sharp contrast to PEP-WT mice, where 20-50% of marginal zone macrophages were LCMV-NP+ (Figure 1F). Thus, despite no detectable difference in viral tropism early after infection between PEP-WT and PEP-619WW mice, there is accelerated clearance of virus from PEP- 619WW mice, which was evident as early as 8 dpi in some cell types.

#### Immune cell heterogeneity in PEP-619WW mice during virus infection

To better understand how the *Ptpn22* alternative allele is impacting anti-viral immunity, we globally characterized all immune cells in the spleen 8dpi (Figure 2A). Live immune cells (CD45+) sorted from 8 dpi infected PEP-WT and PEP- 619WW mice were used for single cell RNA sequencing (scRNAseq) using 10x Genomics platform (Figure 2A). Expression of *Cd45* confirmed most cells sequenced were immune cells (Figure 2B). Using *Csfr1*, *Ly6G*, *Cd19*, and *Cd3e* expression we were able to visualize clustering of monocytes, neutrophils, B cells, and T cells, respectively (Figure 2C). Next, based on these markers we reclustered the specific cell populations to group transcriptionally similar cells and then we compared the proportion of each new cluster within each genotype.

**Figure 2.**
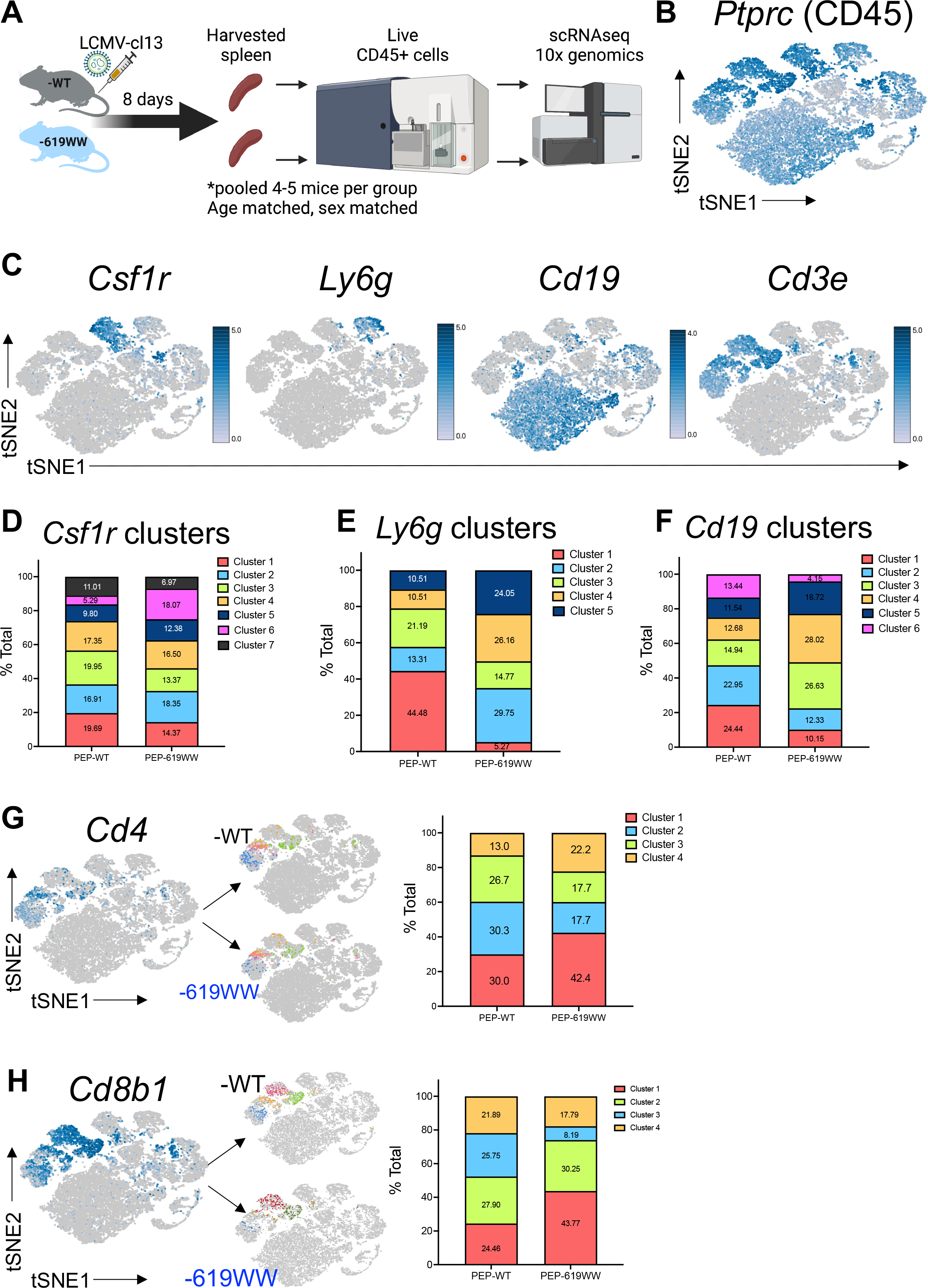
PEP-619WW mice have transcriptionally distinct immune cell subsets from PEP-WT mice during LCMV-cl13 infection. At 8dpi, pooled splenocytes (by genotype) from age matched, sex matched mice were sorted for live CD45+ cells and submitted for single cell RNA sequencing (scRNAseq) on the 10x genomic platform (A). Relative gene expression of *Ptprc* indicating immune cell in aggregated data set (both genotypes) (B). Relative gene expression to located immune cell clusters of myeloid *Csfr1*, neutrophil *Ly6g*, B cell *Cd19,* and T cell *Cd3e* (C). Using Loupe software, each population was reclustered to group like cell subsets among the larger cell type and the proportions compared between genotypes. The frequency of each cluster among total cell population is listed on the graph. Myeloid cell cluster, based on *Csf1r* expression (D), Neutrophil clusters, based on *Ly6g* expression (E), and B cell cluster, based on *Cd19* expression (F). T cells were further broken down into CD4 T cells (expression of *Cd3e*>0, *Mt2*<10, *Cd4*>0) (G) and CD8 T cells (expression of *Cd3e*>0, *Mt2*<10, *Cd8b1*>0) (H). tSNE plots highlighting *Cd4* (G) and *Cd8b1* (H) as well as breakdown of cluster for each T cell type in -WT and - 619WW cells. Corresponding quantification of proportion of either CD4 T cells or CD8 T cell subsets is next to tSNE plot.

Amongst *Csfr1* expressing monocytes 7 unique clusters were identified. Cluster 6 demonstrated the greatest difference in proportion between PEP-WT and PEP- 619WW cells (Figure 2D). *Ly6G* expressing cells were grouped into 5 unique clusters, all of which were represented at different proportions in PEP-WT and PEP-619WW mice (Figure 2E). *Cd19* expressing cells were grouped into 6 unique clusters, the proportions of which also differed between PEP-WT and PEP-619WW mice (Figure 2F). These data show that PEP-WT and PEP-619WW mice have different proportions of monocytes, neutrophils, and B cell subsets present in the spleen at 8 dpi and suggests that the PEP-619WW allele pleiotropically impacts the anti-viral immune response.

#### CD4 T cell transcriptional identity in infected PEP-WT and PEP-619WW mice

It is well established that the clearance of LCMV is dependent on virus specific CD4 and CD8 T cell responses [35, 36]. Therefore, we set out to define how the *Ptpn22* pro-autoimmune allele affected T cell populations during LCMV-cl13 infection. We identified CD4 T cell and CD8 T cells using *Cd3e* expression, in addition to *Cd4* or *Cd8b1* (Figure 2G-H). These populations were then clustered in an unbiased form using Loupe software and we looked at the proportion of each cluster amongst all CD4 or CD8 T cell clusters from PEP-WT or PEP- 619WW mice (Figure 2G-H).

CD4 T cells reclustered into 4 distinct populations, which differed in proportion between PEP-WT and PEP-619WW mice (Figure 2G). Among CD4 T cells in PEP-619WW mice, cluster 1 was most represented, and made up 42.4% of CD4 T cells, whereas it represented only 30% of PEP-WT CD4 T cells (Figure 2G).

Furthermore, we observed decreased proportions of cluster 2 and 3 among PEP- 619WW CD4 T cells compared to PEP-WT CD4 T cells. Cluster 4 had a higher proportion of cells (22.2%) amongst PEP-619WW compared to PEP-WT (13.0%) (Figure 2G).

To learn about the phenotype and function of these CD4 T cell subsets, we looked at the top defining genes for each cluster (Figure 3A; D). Cluster 1 is defined by a set of T follicular helper cell (TFH) genes: *Sostdc1*, *Tbc1d4*, *Izumo1r*, and *2310001h17rik* [37–42]; and activated and effector CD4 T cell genes: *Tnfsf8*, *Rgs10*, *Ypel3*, *Cd200*, and *Malt1* (Figure 3A) [43–49]. Increased expression of both *Cd200* and *Malt1* is associated with increased TCR signaling [47, 49]. This transcriptional data suggests that cluster 1, which is higher in proportion in PEP- 619WW mice, is likely comprised of TFH and activated/effector-type CD4 T cells.

**Figure 3.**
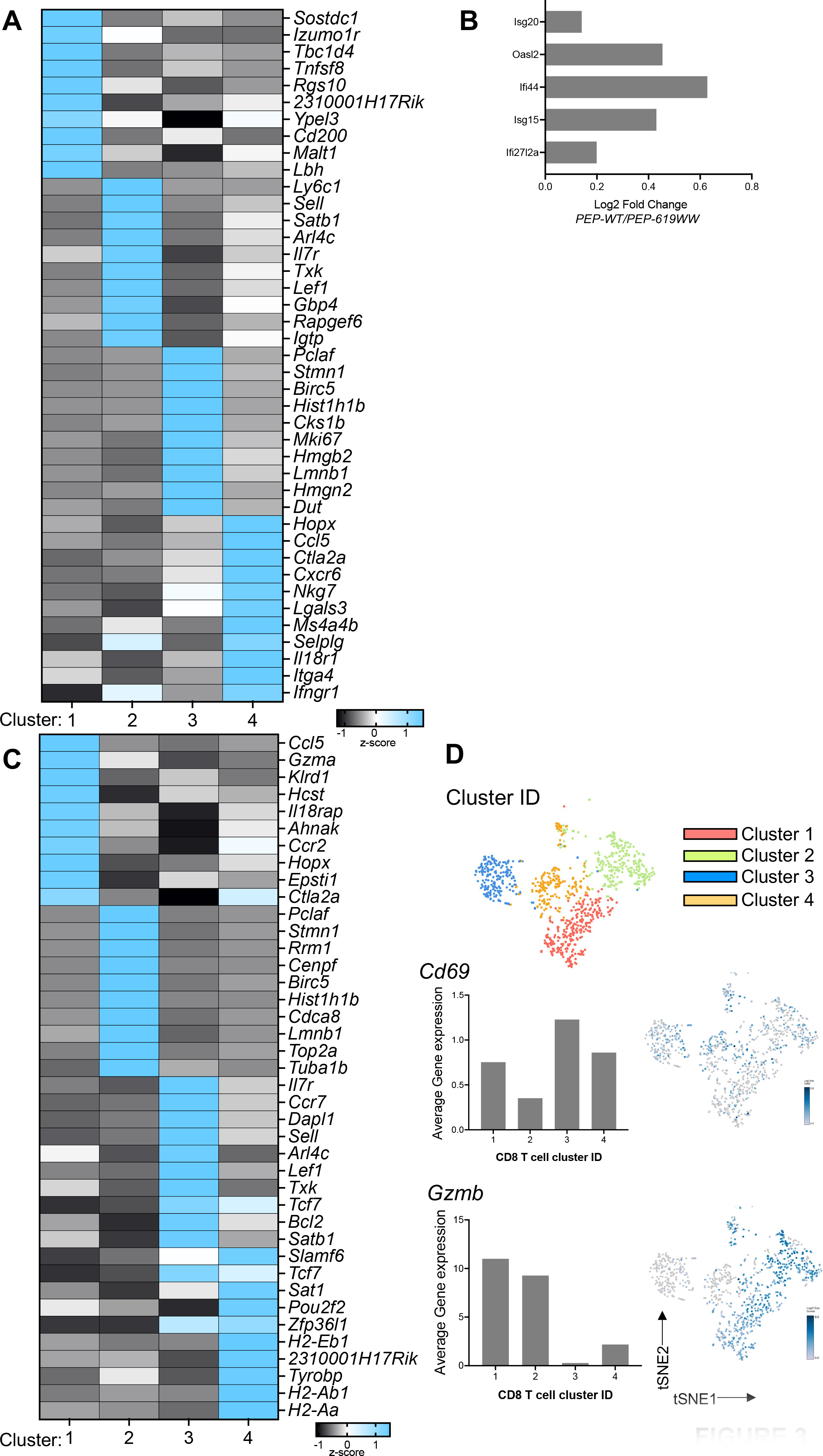
Top unique genes defining CD4 and CD8 T cell clusters from other clusters. Using Loupe, we identified the distinguishing gene between the CD4 T cell cluster (A) or CD8 T cell cluster (C). We then calculated the z-score of these genes across the different cluster populations within each T cell subset (-1 black, to +1 brighter blue). Genes associated with each cluster are listed. Cluster ID on bottom. Log fold change of selected interferon inducible genes, *Isg20, Oasl2, Ifi44, Isg15,* and *Ifi27l2a* (B). Average gene expression of selected genes to further identify CD8 T cell clusters *Cd69* and *Gzmb* quantified on bar graph for each CD8 T cell cluster and highlighted on tSNA visualization (D).

CD4 T cell cluster 2 exhibited high expression of naïve CD4 T cell markers *Arl4c*, *Sell*, *Gpb4*, *Il7r*, and *Rapgef6* [46, 50–53] (Human Protein Atlas) [46]as well as high expression of genes that are key for T cell development and lineage commitment, including *Satb1*, *Txk*, and *Lef1* [54–58] (Figure 3A). Many genes associated with cluster 3 are linked to proliferation and survival, including *Birc5*, *Hist1h1b*, *Cks1b*, *Mki67*, *Hgmn2*, *Lmnb1*, *Hgmn1*, and *Dut* [59–63] (Figure 3A).

Genes upregulated in CD4 T cell cluster 4 are associated with activation (*Hopx*, *Ctla2a*, *Cxcr6*, *MS4ab4*), effector functions (*Ccl5*, *IL18r1*, *Ifngr1*), and migration/homing (*Selplg*, *Itga4*) (Figure 3A) [64–66] [67–73]). Also, CD4 T cell cluster 4 is defined by expression of *Nkg7*, which is associated with cytotoxic CD4 T cell function (Figure 3A) [51]. PEP-619WW also had a higher proportion of cluster 4 cells, which transcriptionally align as cytotoxic or effector CD4 T cells.

Taken together, these data suggest that in response to infection, PEP-619WW mice have more effector-like CD4 T cells (cluster 1 and 4) as compared to PEP-WT mice.

Prolonged type I IFN (IFN-I) during LCMV-cl13 infection is associated with viral persistence [26, 74]. Therefore, we next wanted to know if CD4 T cells from PEP-WT and PEP-619WW had different IFN-induced gene signatures. Using our scRNAseq 10x genomics data we looked at the change of expression in various interferon stimulated genes (ISGs) in PEP-WT and PEP-619WW CD4 T cells.

*Isg20*, *Oasl2*, *Ifi44*, *Isg15*, and *Ifi27l2a* all have a log2 fold change greater than 0, indicating higher relative expression in PEP-WT mice (Figure 3B), suggesting PEP-619WW CD4 T cells had IFN-I signaling cells.

#### CD8 T cell transcriptional identity in infected PEP-WT and PEP-619WW mice

Reclustering of CD8 T cells also grouped the cells into 4 distinct cell populations (Figure 2H). In PEP-WT mice, there was a relatively even distribution of each CD8 T cell cluster. When comparing the proportion of each cluster amongst all CD8 T cells in PEP-619WW mice, 2 of the clusters stood out as interesting based on percent of total CD8 cells within each genotype. Cluster 1 appeared overrepresented in the PEP-619WW (43.77%) mice compared to PEP-WT (24.46%) whereas cluster 3 was less represented in PEP-619WW CD8 T cells (8.19%) compared to PEP-WT CD8 T cells (25.75%) (Figure 2H).

Using their distinct transcriptional signatures, we further analyzed the CD8 T cell clusters of interest. Genes associated with T effector cell phenotype, *Gzma*, *Ccr2*, *Il18rap*, *Ahnak*, *Cxcr6*, and *Ifngr1* are upregulated in cluster 1 (Figure 3C) [39, 75, 76]. Additionally, using iPathway, the top three pathways associated with cluster 1 are viral protein interaction with cytokine and cytokine receptor, cytokine-cytokine receptor interaction, and cell adhesion molecules (Table 1).

**Table 1.**
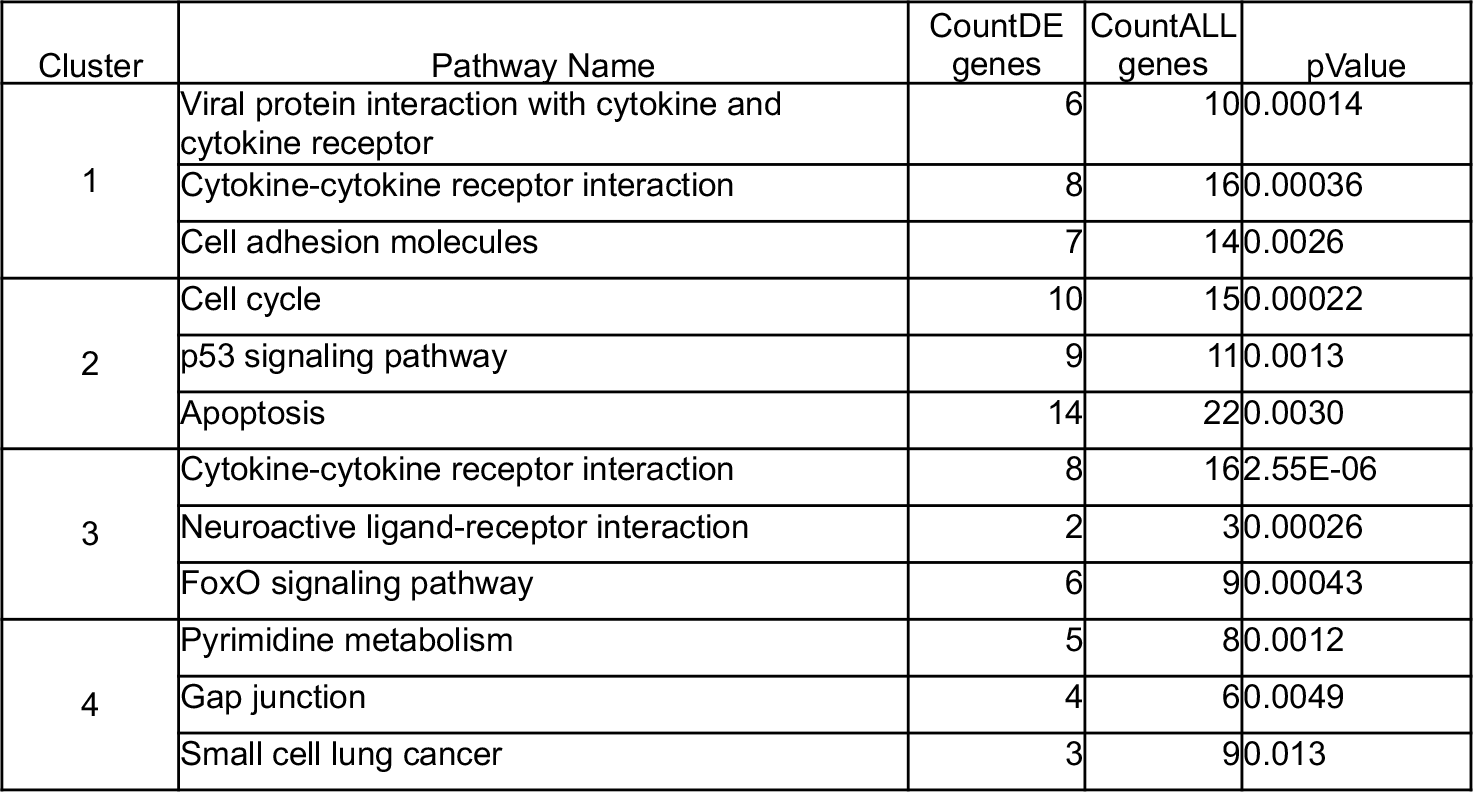
Top Pathways upregulated within each CD8 T cell cluster of LCMV-13 infected mice at 8dpi. Gene expression data exported from Loupe for each CD8 T cell cluster, as identified in Figure 2, was uploaded to iPathway analysis to determine pathways which were significantly associated with those gene sets. Table shows the top 3 pathways for each indicated cluster, the count of differentially expressed (DE) genes that were expressed in each cluster associated with the pathways, count all genes in that pathway, and the p-value associating that pathway with the gene expression data set.

| Cluster | Pathway Name | CountDE genes | CountALL genes | pValue |
| --- | --- | --- | --- | --- |
| 1 | Viral protein interaction with cytokine and cytokine receptor | 6 | 10 | 0.00014 |
|  | Cytokine-cytokine receptor interaction | 8 | 16 | 0.00036 |
|  | Cell adhesion molecules | 7 | 14 | 0.0026 |
| 2 | Cell cycle | 10 | 15 | 0.00022 |
|  | p53 signaling pathway | 9 | 11 | 0.0013 |
|  | Apoptosis | 14 | 22 | 0.0030 |
| 3 | Cytokine-cytokine receptor interaction | 8 | 16 | 2.55E-06 |
|  | Neuroactive ligand-receptor interaction | 2 | 3 | 0.00026 |
|  | FoxO signaling pathway | 6 | 9 | 0.00043 |
| 4 | Pyrimidine metabolism | 5 | 8 | 0.0012 |
|  | Gap junction | 4 | 6 | 0.0049 |
|  | Small cell lung cancer | 3 | 9 | 0.013 |

These pathways are associated in actively responding CD8 T cells. PEP-619WW mice had a higher proportion of cluster 1 cells amongst total CD8 T cells compared to PEP-WT mice, suggesting PEP-619WW mice had more anti-viral effector CD8 T cells.

Cluster 3, which was proportionally more represented in PEP-WT mice, has numerous genes associated with either naïve CD8 T cells, memory CD8 T cells, and the transition from effector to memory cells (Figure 3C). These include *Il7r*, *Ccr7*, *Sell*, *Lef1*, and *Tcf7* [39, 75–77]. Additionally, *Tcf7, Il7r* and *Bcl2* have been implicated in the memory/T cell exhaustion precursor cell population [75, 76]. If cluster 3 was largely composed of activated effector cells, we would expect high expression of *Cd69* along with *Gzmb* (Figure 3D)*. Cd69* is most highly expressed in cluster 3, indicating that this group is largely comprised of antigen experienced T cells. However, cluster 3 had the lowest expression of *Gzmb,* suggesting these cells are not functionally active. Pathways significant in cluster 3 are cytokine- cytokine receptor interaction, neuroactive ligand-receptor interaction, and the FoxO signaling pathway (Table 1). FoxO signaling is known to restrict T cell effector programs and terminal differentiation following LCMV infection [78].

Taken together, the expression of *Tcf7*, *Il7r*, and *Bcl2*, along with expression of *Cd69*, and FoxO signaling suggests that CD8 T cell cluster 3 contains pre- exhausted T cells. Infected PEP-WT animals had a higher proportion of cluster 3 among the total CD8 T cells compared to PEP-619WW mice.

Within CD8 T cell cluster 4, some of the top genes are associated with the exhausted T cell progenitor pool such as *Slamf6*, *Tcf7*, and *Pou2f2* (Figure 3D) [79]. Surprisingly, antigen presentation genes, including *H2-Eb1*, *H2-Ab1*, and *H2-Aa* are also expressed in this cluster. To further confirm that cluster 4 is composed of CD8 T cells, we looked at the relative expression of *Cd3e* and *Cd8b1* in each CD8 T cell cluster (Supp Fig 1A-C). All 4 clusters express these CD8 T cell markers. We also confirmed expression of the class II genes (Supp Fig1D-F). To determine if there were antigen presenting cells (APCs) contaminating our reclustered CD8 T cell population, we also looked at the expression of myeloid and B cell markers, *Lyz2*, *Csfr1*, and *Cd19*. Although there was no expression of *Cd19* in our CD8 T cell cluster, some cells in cluster 4 did express of *Lyz2* and *Csf1r*. The same cells also express *Cd3e* and *Cd8b1*.

Importantly, expression of class II genes does overlap with that of *Lyz2* and *Csf1r*. Although class II expression in human T cells has been reported during Hepatitis C Virus (HCV), HIV-1, Ebola Virus, Dengue Virus, and SARS-CoV2 infections in humans, there is not strong evidence for this same observation in mice [80–85]. Pathways that connected with CD8 T cell cluster 4 are pyrimidine metabolism (p=0.001), gap junction (p=0.004), and small cell lung cancer (p=0.0133) (Table 1). From this data we are unsure as to the identity of this cell population and future studies are needed to determine if class II gene expression in mice during virus infection is biologically relevant or if this is a transcriptional artifact in our study.

#### Pro-autoimmune allele enhances antigen specific T cell activation

##### Anti-viral CD4 T cell function

We next wanted to confirm that the transcriptional difference identified by scRNA-seq resulted in differences in protein expression and T cell function following infection. Th1 polarized CD4 T cells express high levels of the transcription factor Tbet and are a major source of IFNγ during viral infection, a cytokine critical for activating innate and adaptive immunity to clear a virus infection [36, 86].

Following infection, PEP-619WW had a higher frequency of Th1 cells, determined by CD44 and Tbet positive cells (Figure 4A, B). We next measured the amount of Tbet in the Th1 population in PEP-WT and PEP-619WW mice. As anticipated, Tbet expression was highest in Th1 cells from PEP-619WW mice 8 dpi (Figure 4C). Although there is a higher proportion of Th1 cells, it is possible that the functions of these T cell could be tempered by the presence of Tregs.

**Figure 4.**
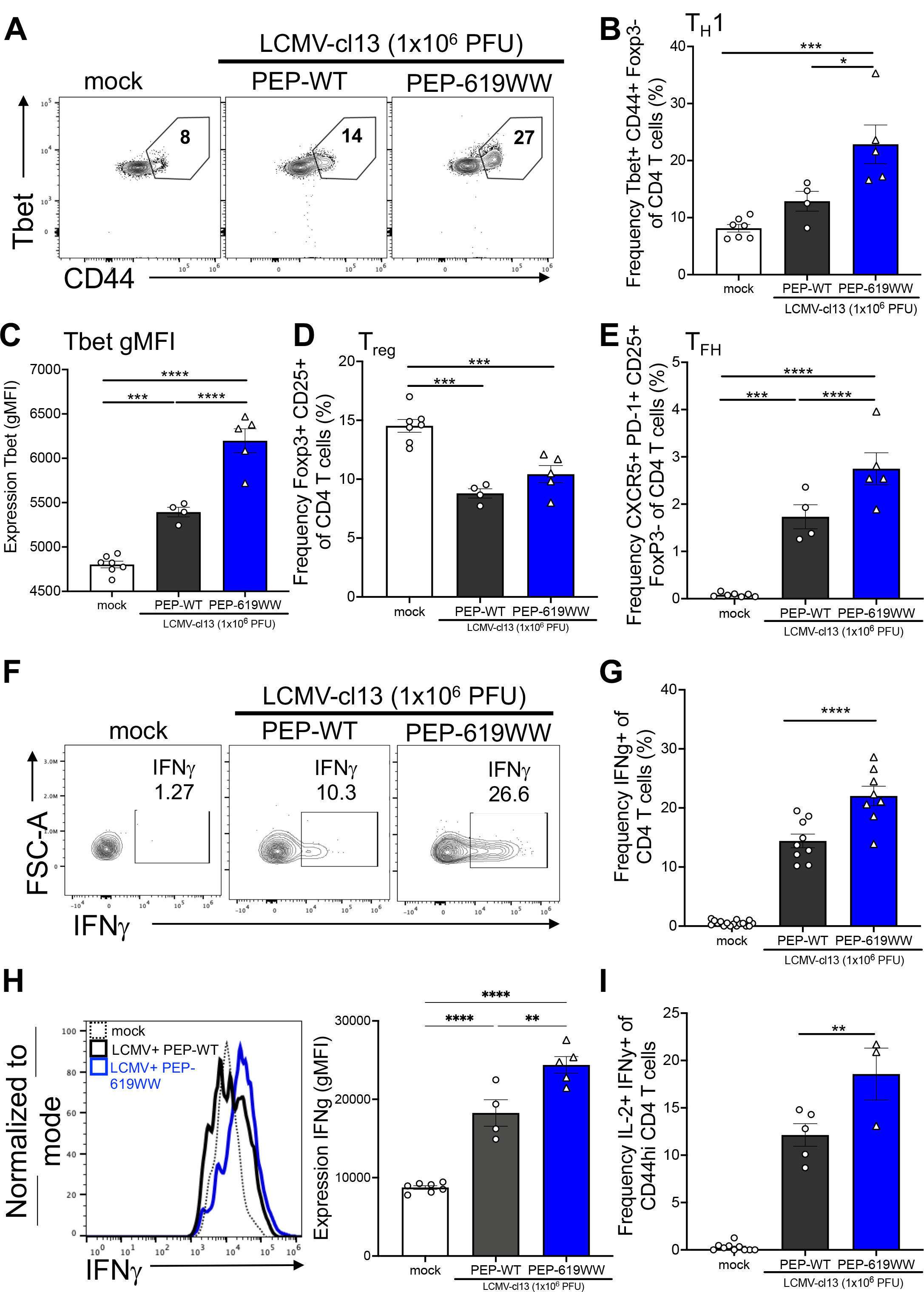
PEP-619WW mice have enhanced CD4 T cell activation during LCMV-cl13 infection. 8dpi splenocytes were isolated and examined for CD4 T cell subsets. Th1 cell (Lymphocytes> single cells x2> live> CD3+ Cd19-> CD4+ CD8α-> Foxp3- CD44+> Tbet+) representative flow plot (A), Th1 frequency of total CD4 (B), and gMFI of Tbet in CD44+ CD4 T cells (C) in mock infected (white bar, circles), infected PEP-WT (black bar, circles), and infected PEP-619WW (blue bar, circles). Frequency of Tregs amongst total CD4 T cells (Lymphocytes> single cells x2> Live> CD19- CD3+> CD4+ CD8α-> Foxp3+ CD25+) (D). Frequency TFH cells (Lymphocytes> single cells x2> Live> CD19- CD3+> CD4+ CD8α-> Foxp3- CD25+> CXCR5+ PD-1+) (E). CD4 T cells were also assessed for IFNγ and IL-2 production after peptide (GP61-80) stimulation (F,G,H). IFNγ+ cells representative flow plot (F). IFNγ+ of CD44+ CD4+ T cells (Lymphocytes> single cells x2> live> CD3+ Cd19-> CD4+ CD8α-> CD44+ >IFNγ+) quantification (G). Expression of IFNγ in CD44+ CD4+ T cells, representative histogram (H) and quantification (H). Frequency of IL-2+ IFNγ+ CD44hi CD4+ T cells (Lymphocytes> single cells x2> live> CD3+ Cd19-> CD4+ CD8α-> CD44+ >IFNγ+ IL-2+). Representative experiment or sample shown for A-F, H-I. Pooled data in G. Each dot represents an individual mouse. Experiments were repeated at least 3 times. SEM shown. *p<0.05, **p<0.01, ***p<0.001, ****p<0.0001. One Way ANOVA with Tukey Post Hoc Analysis.

However, examination of the proportion of Foxp3 expressing CD4 cells in LCMV infected animals showed that both genotypes exhibited a decrease in Tregs following infection (Figure 4D).

To directly test CD4 T cell anti-viral function, whole splenocytes were stimulated with the LCMV CD4 T cell immunodominant epitope peptide (GP61-80) at 8 dpi. In this system, the antigen specific activation of the CD4 T cells reflects their function *in vivo*. A higher proportion of PEP-619WW CD44+ CD4 T cells produced IFNγ in response to stimulation when compared to the cells from PEP-WT infected mice (Figure 4F,G). Also, individual cells from infected PEP-619WW mice had a higher level of IFNγ production (Figure 4H). Furthermore, following peptide stimulation, PEP-619WW mice had a higher proportion of polyfunctional CD4 T cells, producing both IFNγ and IL-2 (Figure 4I). These data demonstrate PEP-619WW mice have a more activated and functional Th1 CD4 T cell population than PEP-WT mice at 8dpi.

Another key function of CD4 T cells during viral infection is differentiation into TFH cells and development of antibody producing B cells in germinal centers, thus contributing to long lasting memory against the viral infection [87]. Of note, PEP- 619WW mice exhibited almost twice the frequency of TFH cells among CD4 T cells, as early as 8 days post infection (Figure 2E). This correlated with increased serum titer of anti-LCMV IgG2A antibody in PEP-619WW mice 9 dpi (Supp Figure 1).

##### Antiviral CD8 T cell function

CD8 T cell function is also required for clearance of LCMV-cl13 [21]. Based on the scRNAseq data and increased function of anti-viral CD4 T cells, we hypothesized that we would see increased CD8 T cell activation and anti-viral function. At 8 dpi, we did not detect a difference in the proportion of activated CD8 T cells between PEP-WT and PEP-619WW mice, as measured by CD44 (Figure 5A). We did observe a slight increase in the frequency of immunodominant epitope, GP33-41 (GP33), specific CD8 T cells (Figure 5B) in infected PEP-619WW mice, compared to infected PEP-WT mice. However, the absolute cell number was not different between genotypes (Figure 5C). When spleen from 8-day infected mice were stimulated with the CD8 T cell immunodominant epitope, GP33, we did not detect any difference in IFNγ production between the two genotypes (Figure 5D,E). These data suggest that at 8 dpi, PEP-619WW does not have a significant impact on CD8 T cell IFNγ production against the immunodominant epitope of LCMV-cl13.

**Figure 5.**
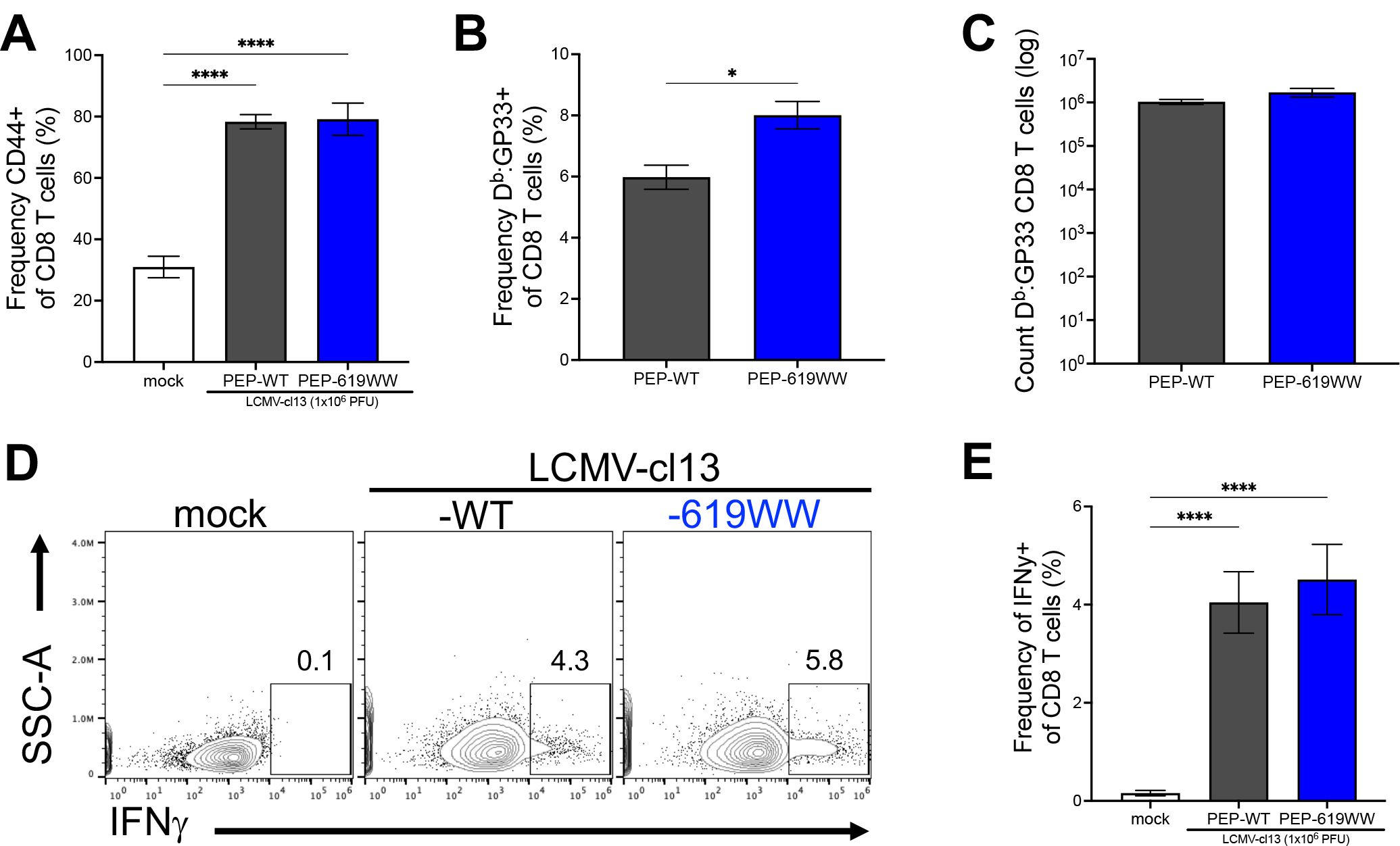
PEP-WT and PEP-619WW have comparable CD8 T cell responses to LCMV-cl13 immunodominant epitope. 8dpi splenocytes from mock infected (white), PEP-WT (black), and PEP-619WW (blue) were isolated and examined for virus specific CD8 T cell subset and assess CD8 T cell function against GP33-41. Frequency of activated CD8 T cells (Lymphocytes> single cells x2> live> CD3+ Cd19-> CD4- CD8α+> CD44+) (A), Db:GP33 tetramer+ CD8 T cells (Lymphocytes> single cells x2> live> CD3+ Cd19-> CD4- CD8α+> Db:GP33+) (B), and absolute number of Db:GP33 CD8 T cells (C). Whole splenocytes were stimulated with GP33-41 peptide. Representative flow plots showing IFNγ production in response to GP33 peptide (D) and quantification (E). Data is pooled form multiple experiments. SEM shown. *p<0.05, **p<0.01, ***p<0.001, ****p<0.0001. One Way ANOVA with Tukey Post Hoc Analysis.

#### PEP-619WW mice have a more immunostimulatory DC phenotype during chronic viral infection

The myeloid compartment contains numerous subsets that may contribute to viral clearance and enhanced CD4 T cell function as observed in PEP-619WW infected mice [25, 32–34, 88]. Classical dendritic cells (cDCs, CD11c+ MHCII+) are critical APCs during viral infection [34]. It is well established that expression of inhibitory ligand PD-L1 suppresses T cell function during LCMV infection [89]. At 8 dpi there is a significant decrease in proportion of splenic PD-L1+ cDCs in PEP-619WW mice (Figure 6 A,B). Furthermore, PD-L1 expression was lower on PEP-619WW cDCs (Figure 6 C,D).

**Figure 6.**
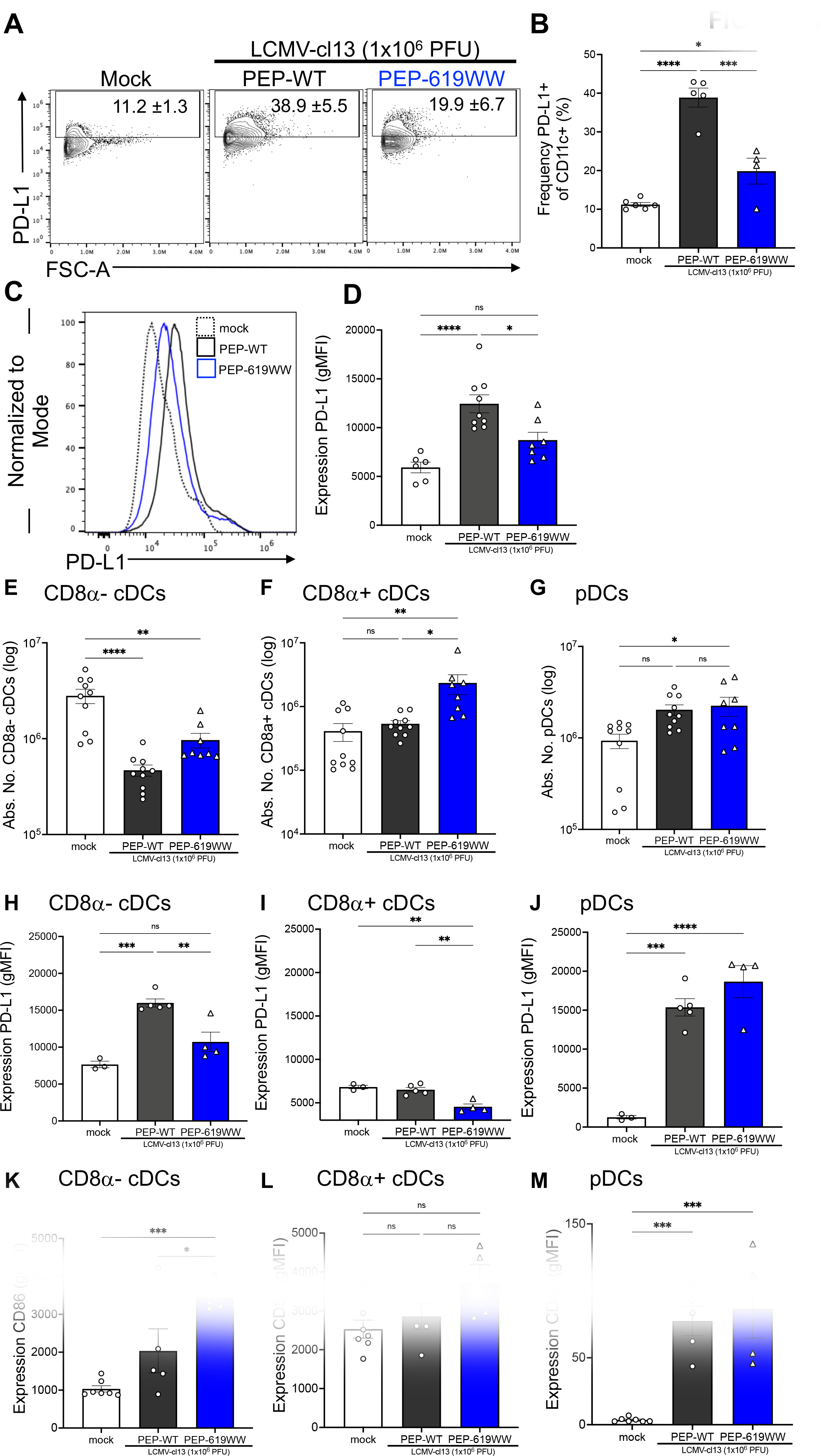
*Ptpn22* pro autoimmune allele promotes more immunostimulatory-like CD8α- cDCs during LCMV-cl13 infection. 8dpi spleens were harvested from mock infected (white, circle), infected PEP-WT (black, circles), and infected PEP-619WW (blue, triangles) mice. Representative flow plot (A) and quantification (B) of PD-L1+ among CD11c+ cells (Lymphocytes> Single cell x2> Live> CD3- CD19-> Ly6G- CD11b+/->CD11c+ F4/80-). PD-L1 expression on CD11c+ cells representative histogram (C) and quantification (D). Absolute number of CD8α- cDCs (Lymphocytes> Single cell x2> Live> CD3- CD19-> Ly6G- CD11b+/->CD11c+ F4/80->CD8α- PDCA-1->MHC-II (I-Ab)+) (E), CD8α+ cDCs (Lymphocytes> Single cell x2> Live> CD3- CD19-> Ly6G- CD11b+/->CD11c+ F4/80->CD8α+ PDCA-1->MHC-II (I-Ab)+) (F), and pDCs ((Lymphocytes> Single cell x2> Live> CD3- CD19-> Ly6G- CD11b+/- >CD11c+ F4/80->CD8α+/- PDCA-1+> Ly6C+ B220+) (G). Expression (gMFI) of PD-L1 on CD8α- cDCs (H), CD8α+ cDCs (I) and pDC (J). Expression (gMFI) of CD86 on CD8α- cDCs (K), CD8α+ cDCs (L), and pDCs (M). Representative experiment or sample shown. Each dot represents an individual mouse. Experiments were repeated at least 3 times. SEM shown. ns=not significant *p<0.05, **p<0.01, ***p<0.001, ****p<0.0001. One Way ANOVA with Tukey Post Hoc Analysis.

Given the diversity of cDC subsets in the spleen, we addressed whether a specific cDC subset had decreased PD-L1 expression or if all DC subsets were affected comparably at 8 dpi. First, we assessed the numbers of different major DC subsets present in PEP-WT and PEP-619WW mice. It has been shown that following LCMV-cl13 infection, there is a significant drop in numbers of cDCs present in the spleen [34, 90]. Indeed, both PEP-WT and PEP-619WW mice showed a significant decrease in the numbers of CD8α^-^ cDCs (CD8α- CD11b+ PDCA-1- CD11c+ DCs) (Figure 6E). However, PEP-619WW mice had significantly more CD8α^+^ cDCs (CD8α+ CD11b- PDCA-1- CD11c+ DCs) at 8dpi compared to PEP-WT mice (Figure 6F). We also looked at plasmacytoid DCs (pDCs) as they are known for their role in supporting clearance of LCMV-cl13 infection [91]. PEP-619WW mice had more pDCs (PDCA-1+ B220+ Ly6C+ CD8α- CD11c+/- DCs) following infection, compared to mock infected animals (Figure 6G). We did not detect a difference in the number of pDCs between mock and infected PEP-WT mice (Figure 6G).

We next examined levels of PD-L1 expression on these same DC subsets at 8dpi. CD8α^-^ cDCs, but not CD8α^+^ cDCs, upregulate PD-L1 in infected PEP-WT mice (Figure 6H,I,J). PEP-619WW CD8α^-^ cDCs have less PD-L1 expression than PEP-WT CD8α^-^ cDCs at 8dpi (Figure 6H). Also, PEP-619WW CD8α^+^ cDCs have less PD-L1 expression compared to infected PEP-WT mice and mock-infected animals. pDCs from both PEP-WT and PEP-619WW have significantly higher levels of PD-L1 compared to uninfected controls (Figure 6J). We also looked at CD86 expression, a receptor that positively regulates T cell activation by DCs. CD86 expression was increased in CD8α^-^ and pDCs in both PEP-WT and PEP-619WW mice following infection. However, the level of increase was significantly greater on CD8α^-^ cDCs from PEP-619WW mice (Figure 6K-M).

Thus, infected PEP-619WW mice have cDC populations with lower PD-L1 expression and enhanced CD86 expression, suggesting a more immunostimulatory phenotype.

#### Ptpn22 pro-autoimmune allele enhances T cell activation through T cell intrinsic and extrinsic mechanisms

We next wanted to know if the enhanced anti-viral CD4 T cell function observed was due to a T cell intrinsic mechanism, such as enhanced TCR signaling, or a T cell extrinsic mechanism, such as a more immunostimulatory cDC phenotype.

Employing adoptive transfer techniques, we constructed mice in which the virus specific CD4 T cells (SMARTA CD4 T cells) expressed an allelic difference in *Ptpn22* (Figure 7A). Post infection, both transferred SMARTA PEP-WT and SMARTA PEP-619WW CD4 T cells expanded, regardless of host genotype (Figure 7 B,C). In the PEP-WT host, SMARTA PEP-619WW CD4 T cells had a higher proportion of IFNγ+ cells than SMARTA PEP-WT CD4 T cells (Figure 7D,E). When SMARTA PEP-WT CD4 T cells were transferred into a PEP- 619WW host, there was a slight increase in the proportion of IFNγ+ virus specific CD4 T cells compared to SMARTA PEP-WT T cells into a PEP-WT host (Figure 7D,E). However, SMARTA PEP-619WW CD4 T cells in a PEP-619WW host had the highest frequency of IFNγ+ virus specific cells (Figure 7D,E). Additionally, SMARTA PEP-619WW cells in a PEP-619WW had the highest expression of IFNγ+ within activated CD4 T cells (Figure 7F). Also, SMARTA PEP-619WW CD4 T cells in a PEP-619WW host had the highest proportion of polyfunctional cells (Figure 7G). Taken together, this suggests the presence of the *Ptpn22* minor allele in either the host or SMARTA CD4 T cells slightly increased T cell function. However, the T cell intrinsic and extrinsic effects have an additive effect on virus specific T cell activation, resulting in the highest amount of IFNγ and IL-2 production.

**Figure 7.**
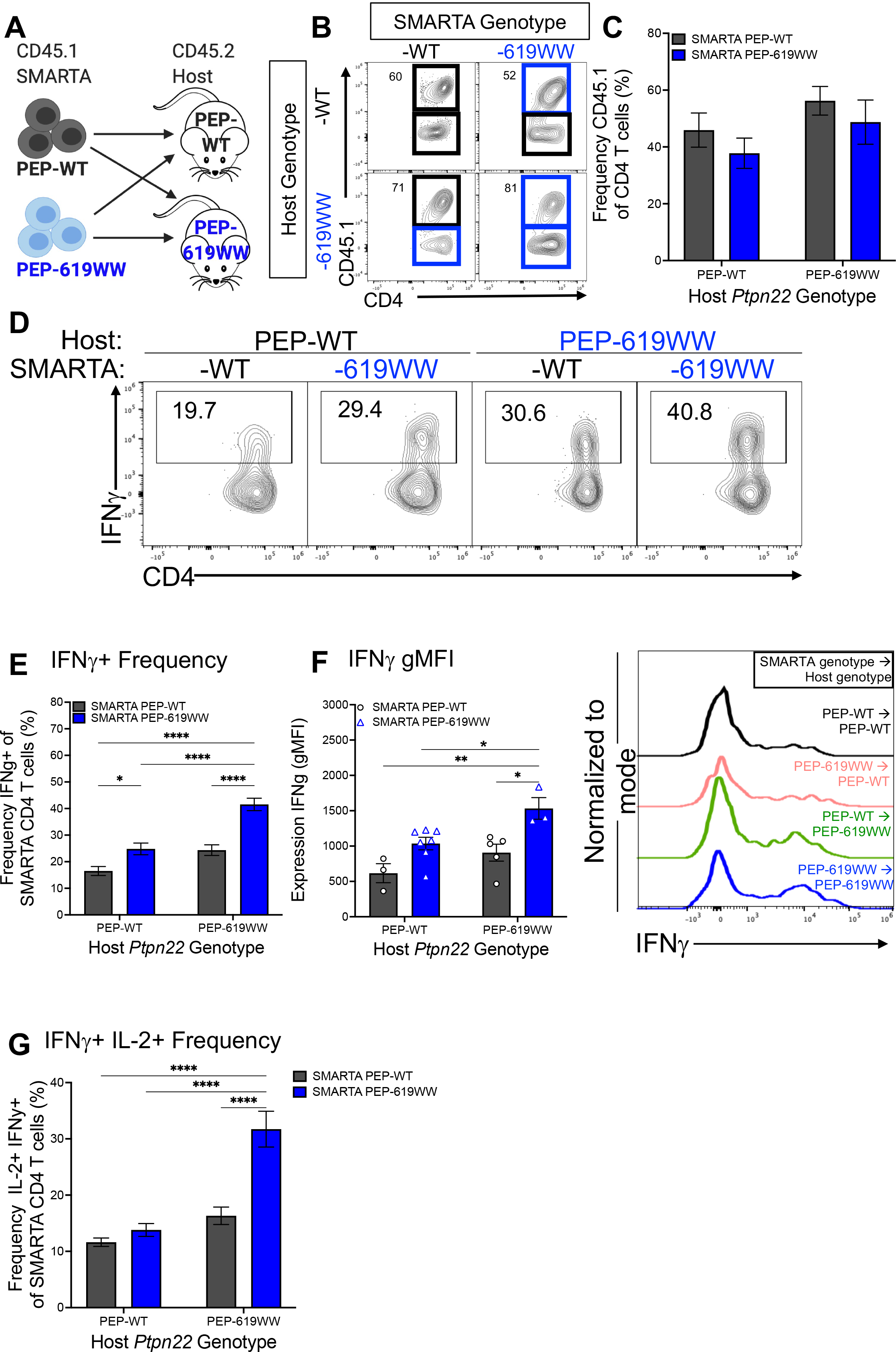
*Ptpn22* alternative allele enhances anti-viral T cell activation through T cell intrinsic and extrinsic mechanisms. CD45.2 C57BL/6 mice (PEP-WT or PEP-619WW) received 5x10^4 CD45.1 SMARTA CD4 T cells prior to being infected with LCMV-cl13 (A). Frequency of CD45.1 SMARTA T cells (Lymphocytes> single cells x2> Live> CD3+ CD19-> CD4+ CD8α-> CD44+> CD45.1+ CD45.2-) for each transfer condition, representative flow plot (B) and quantified (C). Following peptide stimulation with GP61-80 peptide, IFNγ and IL-2 production of SMARTA cells were measured (D-G). Representative flow plot showing IFNγ+ CD45.1+ CD4 T cells (D), quantification IFNγ+ CD45.1 Cd4 T cells (E), expression (gMFI) of IFNγ in CD45.1 SMARTA cells with representative histogram (F), frequency of IFNγ+ IL-2+ CD45.1 CD4 T cells (G). C, E, and G from pooled data. F is a representative experiment. Experiments were repeated at least 3 times. SEM shown. *p<0.05, **p<0.01, ***p<0.001, ****p<0.0001, Two Way ANOVA with Sidak post hoc analysis.

## Discussion

The strong genetic link between the *PTPN22* minor allele (R620W) and autoimmunity would be expected to be an evolutionary disadvantage [7]. However, an advantage in eradicating pathogens could help explain why the variant persists in the population [92, 93]. Thus, it was of interest to assess the impact of the *PTPN22* autoimmunity associated allele has during virus infection. Previously, using mice deficient in *Ptpn22* expression (PEP-null), we demonstrated that *Ptpn22* promotes T cell exhaustion, thus permitting LCMV- cl13 persistence [28]. Although some studies have found that the lack of *Ptpn22* is similar in phenotype to R620W (R619W in mice) several differences have also emerged. This may be because the 620W (619W) variant does not directly affect the phosphatase activity of this enzyme [14, 94, 95]. Rather, the presence of tryptophan disrupts the ability of the enzyme to bind with other proteins such as CSK and TRAF3 [13, 15, 95]. As an example of differing consequences, whereas PEP-null mice exhibit an increased frequency and potency of Tregs as compared with WT [96], this same Treg phenotype is not observed in naïve mice bearing the *Ptpn22* pro-autoimmune allele [19, 97]. Despite its importance in human health, studies have not investigated whether the *Ptpn22* autoimmunity associated minor allele impact anti-viral immune responses.

Using the well-established model of LCMV-cl13, we tested the ability PEP- 619WW mice to overcome a potentially chronic viral infection. PEP-619WW mice largely cleared virus infection and have reduced weight loss compared to wildtype (PEP-WT) mice (Figure 1 A-D). Considering comparable LCMV-NP expression at 3 dpi, our data suggests that clearance is not likely due to early changes in tropism (Figure 1E). By day 8 PEP-619WW demonstrated earlier viral clearance in monocytes, marginal zone macrophages, and pDCs, but this does not correlate with reduced sera titer (Figure 1F). Future studies will address the mechanism(s) in PEP-619WW mice driving viral clearance from these myeloid subsets. Taken together, these data demonstrate expression of PEP-619WW, rather than the WT allele, confers strong protection from virus persistence.

Failure to clear LCMV-cl13 infection is associated with immune cell dysfunction [26, 27, 34, 74, 98]. The use of scRNAseq allowed us to transcriptionally define and interrogate multiple immune cell types in both PEP-WT and PEP-619WW mice during LCMV-cl13 infection (Figure 2). These data suggest the impact of PEP-619WW during viral infection is pleiotropic and impacts multiple mechanisms that result in altered B cell, myeloid cell, and T cell responses. The details of how PEP-619WW impacts different molecular mechanisms within each cell during virus infection is of ongoing interest.

Both CD4 and CD8 T cells are necessary to clear LCMV-cl13 [27]. We investigated the T cell compartment from infected PEP WT and PEP-619WW mice at a transcriptional and functional level. Anti-viral CD4 T cells, but not CD8 T cells, were functionally different in infected PEP-619WW mice. Among CD4 T cells, PEP-619WW mice had higher frequencies of TFH and activated/effector-type CD4 T cells during infection than PEP-WT animals (Figure 3A, Figure 4E). During chronic viral infection, sustained TCR signaling is shown to lead to more TFH cell differentiation compared to acute viral infection [98]. Since *Ptpn22* tempers TCR signaling, disrupting that regulatory mechanism, through presence of the allelic variant (PEP-619WW) may be what is driving this TFH increase during infection. It has been shown that *Ptpn22* pro-autoimmune allele bearing mice and *Ptpn22* knock out mice have increased TFH and germinal center formation [96, 99]. We also show that transcriptionally and functionally, PEP- 619WW have more activated and anti-viral effector CD4 T cells. CD4 T cells from PEP-619WW infected mice produced more IFNγ, and IL-2 compared to PEP-WT cells (Figure 4F-I). Presence of these polyfunctional cells has been linked to be better anti-viral immune response. Taken together, our data demonstrate PEP- 619WW have enhanced anti-viral CD4 T cell responses.

Both T cell intrinsic and extrinsic mechanisms could explain the enhanced anti- viral CD4 T cell function in PEP-619WW mice. Modifying TCR strength through extrinsic signals, such as increased immunostimulatory molecules, or intrinsic signals have an impact on downstream effector functions like IFNγ production [100, 101]. Using murine models, human T cell culture systems, and clinical data researchers have defined a role of *PTPN22/Ptpn22* in tempering the TCR signaling pathway. In these systems, researchers have also shown that the pro-autoimmune allele of *PTPN22* results in sustained TCR signaling, leading to increased T cell activation [14, 94]. Using adoptive transfers studies, we concluded that neither a PEP-619WW T cell nor host alone is sufficient to account for the heightened T cell activation during LCMV-cl13 infection. Rather, there is an additive effect between the PEP-619WW T cell, and PEP-619WW non-T cells resulting in increased CD4 T cell activation. Taken together with our cDC profiling data, it is likely that the large increase of CD4 T cell activation during LCMV-cl13 infection is from additive effects of the CD8α^-^ cDC being a more activator-like phenotype and the CD4 T cell having intrinsic properties to increase T cell activation.

Our data suggest that the presence of the *Ptpn22* autoimmunity associated allele affects multiple parts of the anti-viral immune response critical for controlling a chronic infection. When the PEP-619WW cDCs prime and activate CD4 T cells, the T cells are interacting with a cDC that expresses less PD-L1 and increased CD86, in turn resulting in more T cell activation. PEP-619WW bearing T cells also demonstrate enhanced T cell intrinsic activation and production of IFNγ, potentially through the decreased tempering of TCR signaling or changed IFN-I signaling [14, 94, 102]. The PEP-619WW DC and PEP-619WW CD4 T cell have an additive effect on anti-viral T cell function, which in turn controls the chronic viral infection. Although we did not observe increased numbers of GP33-specific CD8 T cells, these data do not rule out the possibility CD8 T cells against other epitopes contribute to viral clearance in PEP-619WW mice. Additional studies isolating the cell autonomous impact of PEP-619WW are needed to determine which cell type is necessary and/or sufficient to cause enhanced anti-viral T cell function and an enhanced anti-viral immune response. Results of this study provide a platform to further investigate the inter-relationships between immune cells to achieve a desired phenotype as well as determine if/how other autoimmunity associated alleles affect the anti-viral immune response.

## Methods

### Mice

Both males and females ranging from 6-12 weeks of age were used in this study. Animals were housed in general housing conditions at Scripps Research. We do not observe any sex-based difference in these studies with our animals. All animal studies were reviewed and approved by Scripps Research Institutional Animal Care and Use Committee (protocol number: 06-0291). C57BL/6 WT mice were originally purchased from Jackson labs, and then bred and maintained in Scripps Animal Facility. *Ptpn22* R619W (PEP-619WW) mice were generated using CRISPR/Cas9 technology on a C57BL/6 background using methods previously reported ([97, 103] in the Mowen Lab. In short, four nucleotides were replaced on exon 14 of *Ptpn22* to insert the BspEI restriction site and cause an arginine (R) to tryptophan (W) amino acid substitution at amino acid position 619. Genotypes were confirmed through PCR using the following primers which flank the mutated region of *Ptpn22*: Forward-5’ AGCTGATGAAAATGTCCTATTGTGA 3’ and Reverse-5’ GTCCCACTGCATTCTGGTGA 3’. After amplification, PCR products are digested overnight at 37°C with the restriction enzyme, BspEI, which is unique to mutated mice. Digested PCR products are run on a 2% agarose gel to visualize digested bands.

For transfer experiments, naïve CD4 T cells were isolated from the spleens of naïve CD45.1 SMARTA PEP-WT, PEP-null, or PEP-619WW C57BL/6 mice using a negative isolation kit (Stem Cell Technologies, Vancouver, British Columbia, Canada). A low number of isolated CD4 T cells, as indicated in figure legend, were transferred into naïve, sex-matched mice intravenously. 1-2 days following cell transfer, recipient mice were infected with LCMV-cl13, or sterile HBSS for mock control.

For infection, mice received 1x10^6^ PFU LCMV-cl13 resuspended in 100uL sterile PBS intravenously (i.v). Mice receiving mock infection received 100uL sterile PBS (i.v). Mock group of mice included both PEP-WT and PEP-619WW genotypes. Mice were removed from study if weight loss was greater than 25% original starting weight.

### Flow Cytometry and Antibodies

Samples were excised and placed into HBSS with 2% FBS. Samples were minced and incubated with Stem Cell Spleen dissociation media (Stem Cell Technologies, Vancouver, British Columbia, Canada) according to manufacturer’s instructions. Following incubation, minced sample was smashed and filtered through 40uM filter to create a single cell suspension. Single cell suspension was counted and resuspended to desired concentration (dependent on experiment) in HBSS with 2% FBS. Single cell suspensions were used for staining and flow cytometric analysis. Cells were stained in serum free HBSS.

All flow cytometry was completed on a spectral cytometer the Cytek Aurora with a 4 laser or 5 laser system (405nm, 488nm, 640nm, 561nm, and 355nm (5 laser only)). Single color stain OneComp eBeads (Thermo Fisher) were used for unmixing. Unmixed files were analyzed using FlowJo Software (BD Biosciences, San Diego, California). Antibodies used in various combinations (depending on experiment) are as follows: Ghost Viability Dye (v510, Tonbo Biosciences, 1:1000 dilution), CD3e (PE-Cy5/AF532, Tonbo Bioscience/Thermo Fischer, 1:200, clone 145-2C11), CD4 (PerCP/BV605, Tonbo Bioscience/Biolegend, 1:200, clone RM4-5), CD8α (APC-Cy7/APC-H7/APC, BD Biosciences, 1:200, clone 53-6.7), CD11c (PE-Cy5.5, Thermo Fisher, 1:100, clone N418), CD11b (PerCP-Cy5.5, Biolegend, 1:200, clone M1/70), F4/80 (Pacific Orange, Thermo Fisher, 1:100, clone BM8), IFNγ (AF647, Biolegend, 1:100, clone XMG1.2), TNFα (PE-Cy7, Beiolegend, 1:100, clone MP6-XT22), IL-2 (PE/BV421, Biolegend, 1:50, clone JES6-5H4), PDCA-1 (Pacific Blue, Biolegend, 1:200, clone 129C1), CD80 (BV421, 1:200, clone 16-10A1), CD86 (BV605, Biolegend, 1:200, clone GL1), PD-1 (PE-Cy7, Tonbo Biosciences, 1:200, clone J43.1), PD- L1 (PE/BV711, Tonbo Biosciences/Biolegend, 1:100, clone 10F.9G2), CD44 (AF700, Biolegend, 1:200, clone IM7), CD62L (FITC, Tonbo Biosciences, 1:100, MEL-14), Ly6C (BV785, Biolegend, 1:200, clone HK1.4), Ly6G (PE-eFlour610, Invitrogen, 1:200, clone IA8), CD206 (AF647, Biolegend, 1:100, clone CO68C2), CD209b (APC, Tonbo Biosciences, 1:200, clone 22D1), NK1.1 (FITC, Biolegend, 1:100, clone PK136), CD19 (BV711, Biolegend, 1:400, clone 6D5), B220 (APC-Cy5.5, Invitrogen, 1:200, clone RA3-6B2), MHC II I-Ab (FITC, Biolegend, 1:200, clone AF6-120.1), CXCR5 (BV605, Biolegend, 1:100, clone L138D7), Tbet (APC/AF647, Biolegend, 1:200), Foxp3 (PE, Invitrogen, 1:100, clone FJK-16s), LCMV-NP (self-conjugated to AF488 using Thermo Fisher AF488 labeling kit per manufacturer’s instructions, BioXcell, 1:50, clone VL-4).

Tetramer staining occurred in the following conditions with HBSS. CD4 tetramer (I-Ab:GP66-77 (1:200)) was stained at 37C for 2 hours, in dark, alone. CD8 tetramer (D^b^:GP33-41 (1:500)) was stained at room temperature, 1 hour, in dark, alone. Following tetramer stains, cell were washed and stained for other surface markers.

All non-tetramer surface markers were stained in HBSS, at 4C, in dark. If intracellular staining for transcription factors was required, Tonbo Foxp3 Fix/Perm kit was used per manufacturer’s instructions. For intracellular cytokine staining BD Cytofix and Permwash was used according to manufacturer’s instructions.

### Virus

LCMV-cl13 generation, tittering, and infection is as previously described [26, 28, 104]. In short, mice were infected with 1x10^6^ PFU LCMV-cl13. Throughout infection, mice were monitored and then euthanized at designated time points. To determine viral load in animals, serum and organs were harvested at designated time points and tittered on Vero cells and calculated as previously described [26].

### Antigen-Specific T cell activation

Splenocytes from LCMV-cl13 or mock infected animals were isolated at designated time points. After red blood cell lysis and filtering through 70uM filter, splenocytes were resuspended in T stimulation media (RPMI containing 10% FBS, 1% of each of the following Penicillin/Streptomycin, HEPES, Sodium Pyruvate, L-Glutamine, Non-Essential Amino Acids, and 55μM 2- Mercaptoethanol). 2x10^6^ splenocytes were plated into 96-U bottom wells in the presence or absence of 5μg/mL GP61-80 peptide (CD4 T cells) or 2μg/mL GP3361- 80 (CD8 T cell) for 1 hour at 37C. After 1 hour incubation with peptide, Brefeldin A (0.4ug/mL) was added and incubated for an additional 4 hours at 37C. Following stimulation and incubation, cell were spun down, washed with HBSS, and proceeded to extracellular and intracellular staining.

### Single Cell RNA Sequencing and Analysis

At 8dpi, spleens from 4-5 age matched, sex matched PEP-WT and PEP-619WW mice were removed, processed (as described above), and resuspended in a single cell suspension. Splenocytes from each genotype were pooled and FACS sorted for Live CD45+ cells. Sorted cells were submitted to the Scripps Research Genome Sequencing Core for single cell RNA sequencing via 10x Genomics platform.

We performed primary and secondary analysis of scRNA-Seq data using CellRanger, an analysis suite consisting of multiple pipelines for end-to-end analysis of single cell data. It uses a custom-built wrapper around Illumina’s bcl2fastq to demultiplex raw base calls. This is followed by removal of duplicates using UMI (unique molecular identifier) counting. These preprocessed samples are aligned using STAR [105], which performs splicing-aware alignment of reads to the genome. Aligned reads are then bucketed into exonic and intronic regions using a reference GTF (gene transfer format) file. Finally, read counts per gene are calculated, and these values are used for downstream estimation of differentially expressed genes. The unbiased single-cell profiles and high- resolution data allows counting and clustering of cells with similar transcriptome profiles to uncover distinct cell subsets and genes preferentially expressed by the cells. We performed dimensionality reduction, clustering, generation of open- standard file formats for interactive analysis, using both: currently most widely adopted methods and algorithms [106] [107] [108] as well as most recent and better performing methods and algorithms [109] [110] [111] now available. We performed interactive exploration and functional analysis, to uncover cells sub- populations, gene markers and differential expression between sample groups, and suggest possible pathways and mechanisms, leveraging leading edge platforms for single cell exploration [112] and complex functional analysis [113].

Visualization, clustering, and population identification, and gene expression was all completed with Loupe Software. t-distributed stochastic neighbor embedding (tSNE) visualization and clustering were performed using aggregated file containing both PEP-WT and PEP-619WW samples.

### Anti-LCMV neutralizing antibody assay

Serum antibody enzyme-linked immunosorbent assays were performed as previously described [114]. Briefly, microplates were coated with LCMV-infected baby hamster kidney (BHK) cell lysates overnight. Subsequently, nonspecific binding was blocked by coating microplates with 3% bovine serum albumin (BSA) in phosphate-buffered saline (PBS). Serial dilutions of serum were carried out in 1% BSA in PBS. After overnight incubation, plates were incubated with purified biotin-conjugated anti-mouse IgG, IgG1, or IgG2a (1030-08, 1070-08, 1080-08; Southern Biotech) antibodies for 2 hrs. Antibody detection was further performed using streptavidin–alkaline phosphatase (11089161001, Roche) for 1 hour and then alkaline phosphatase substrate solution containing 4-nitro-phenyl phosphate (N2765-50TAB, Sigma-Aldrich) for approximately 30min. A CLARIOstar Plus microplate reader was used to quantify the results.

### Statistical Analysis and Graphing

All statistical analysis were performed using GraphPad Prism (La Jolla, CA) and used as appropriate for the data. The type of statistical test is listed in figure legends. Data was considered statistically significant is the p value<0.05.

Graphs were made in GraphPad Prism (La Jolla, CA). Figure legends indicate if data shown is pooled from multiple studies or is from representative study.

### Data Availability

Any data will be made available upon request. All scRNA sequencing data is being uploaded. Prior to publication we will supply the GEO accession number for the data sets.

## Author contributions

R.C.O. conceptualized, designed, and performed experiments; analyzed results; reviewed data and wrote and edited the manuscript. K. Marquardt provided reagents, bred and genotyped mice, and performed experiments. I.P. performed experiments. K. Mowen was the principal investigator whose laboratory generated the PEP-619WW mice on the C57BL/6 background. A.D. assisted in single cell RNA sequencing processing and analysis. J.R.T. conceptualized and designed experiments, reviewed data, edited manuscript. L.A.S. conceptualized and designed experiments, reviewed data, and wrote and edited the manuscript.

## Acknowledgements

We thank Scripps Research Flow Cytometry Core, Histology and Microscopy Core, and Vivarium staff for their expertise and assistance in this work. We also thank T. Fehr and C. Antunes (University of Kansas) for their careful reading and editing of this manuscript. Since the creation of the PEP-619WW mice, K. Mowen has passed away and is greatly missed. The authors declare no competing financial interests.

## Abbreviations

T1D: Type I Diabetes
Lyp: Lymphoid protein
PEP: PEST-domain Enriched Phosphatase
TCR: T Cell Receptor
BCR: B Cell Receptor
TRAF3: TNF receptor associated factor 3
CSK: C-terminal Src Kinase
TLR: Toll-Like Receptor
LCMV-cl13: Lymphocytic choriomeningitis virus clone 13
Treg: T regulatory cell
Dpi: days post infection
LCMV-NP: Lymphocytic choriomeningitis virus Nucleoprotein
cDC: conventional Dendritic Cell
pDC: plasmacytoid Dendritic Cell
IFN-I: Type I Interferon
IFNAR1/2: Interferon alpha receptor 1 and 2
ISG: Interferon stimulated gene
Th1: T Helper 1 cell
T_FH_: T Follicular helper cell
IFNγ: Interferon gamma
TNFα: Tumor necrosis factor alpha
IL-2: Interleukin 2
GP: glycoprotein
PD-L1: Program death receptor ligand 1
WT: Wildtype
APC: antigen presenting cell

---

**Supplementary Figure 1.**
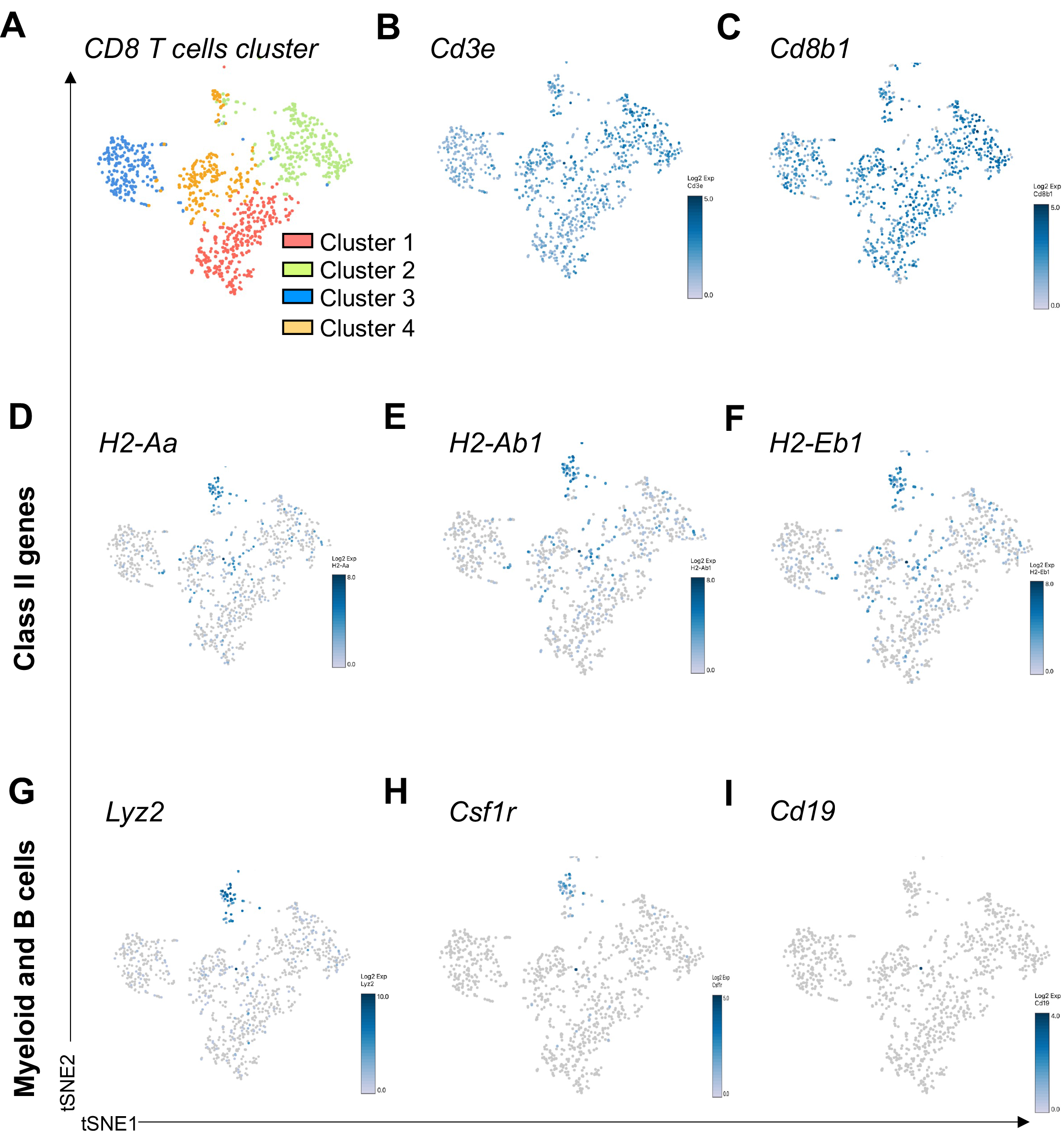
Gene expression in CD8 T cell recluster. CD8 T cells were reclustered within Loupe software using aggregated file containing both PEP-WT and PEP-619WW data. tSNE visualizing Clusters 1-4 of CD8 T cell re-cluster analysis. Cluster 1 salmon, Cluster 2 light green, cluster 3 blue, and Cluster 4 light orange (A). Average gene expression of genes *Cd3e* (B), *Cd8b1* (C), *H2-Aa* (D), *H2-Ab1* (E), *H2-Eb1* (F), *Lyz2* (G), *Csfr1* (H) and *Cd19* (I). Gene expression shown in Log2 scale, range is on heat map scale for each tSNE plot.

**Supplementary Figure 2.**
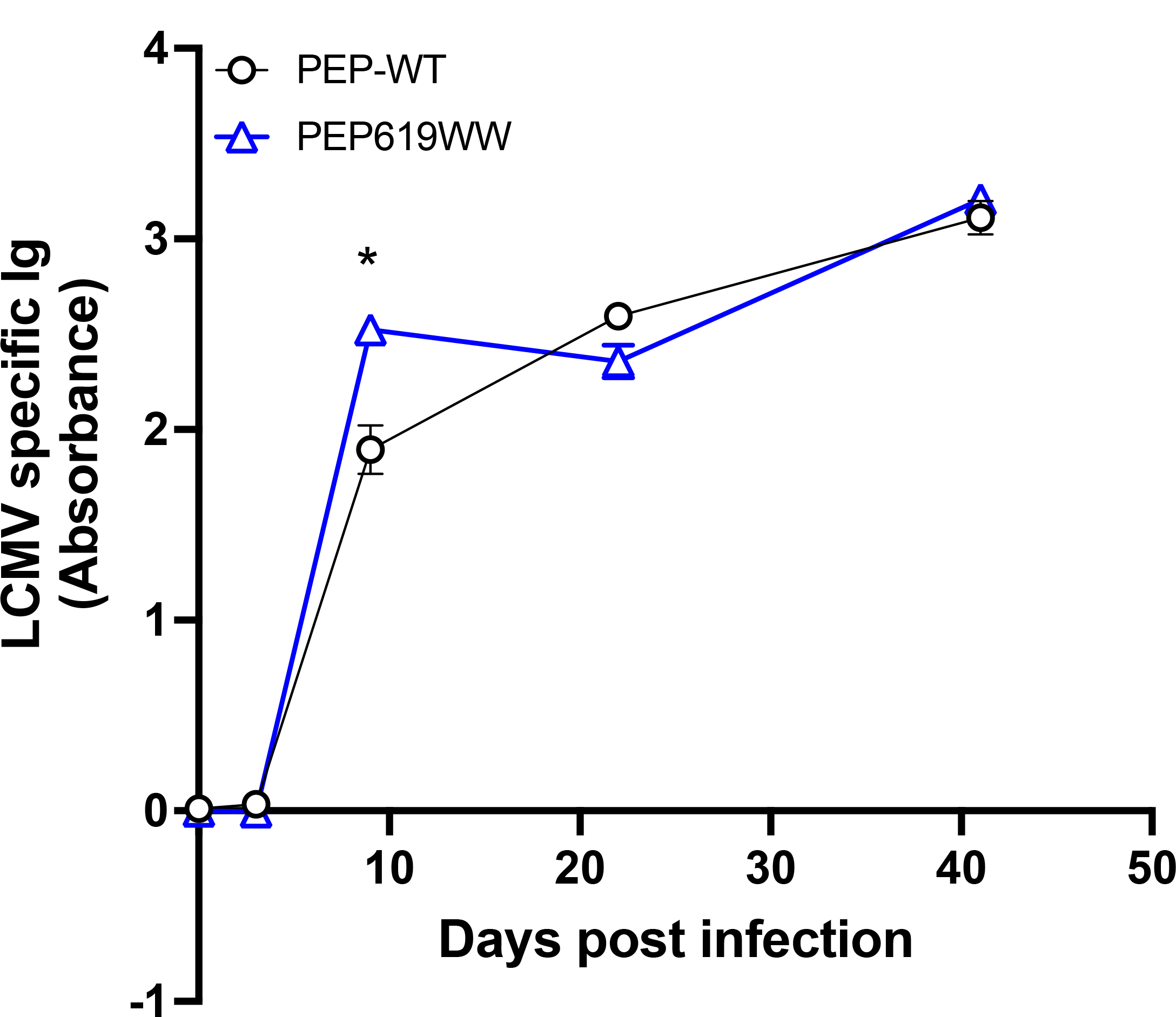
PEP-619WW have more LCMV neutralizing antibodies 9 days post infection than PEP-WT. Serum from infected PEP-WT (black) and PEP-619WW (blue) taken at 0, 3, 9, 21, and 41 days post infection and measured for neutralizing IgG2A antibodies. Data shown from a 1: 1583 dilution. Representative experiment shown. SEM shown. *p<0.05; T-test at each time point between genotypes.

